## Supplemental Data 1 for "Interdependent progression of bidirectional sister replisomes in *E. coli*": Chen supplemental biorxiv.pdf

**Supplementary table 1.** Bacterial strains and plasmids used in this study.

| Strain | Genotype or description | Source |
|---|---|---|
| DB1568 | FRT::*dnaN-ypet* | This study |
| DB2146 | 141x *tetO* array @ *lac* (+1.10 Mb), pDM15 | (Joshi *et al.*, 2011) |
| DB2185 | 141x *tetO* array @ *gln* (+0.13 Mb), pDM15 | (Joshi *et al.*, 2011) |
| DB2193 | 141x *tetO* array @ *dnaB* (+0.34 Mb), pDM15 | (Joshi *et al.*, 2011) |
| DB2260 | Δ*recB745*::FRT*kan*FRT | JW 2788 (Baba *et al.*, 2006) |
| DB2403 | 141x *tetO* array @ *gln* (+0.13 Mb), Δ*recB745*::FRT*kan*FRT, pDM15 | This study |
| DB2722 | 141x *tetO* array @ *dnaB* (+0.34 Mb), Δ*recB745*::FRT*kan*FRT, pDM15 | This study |
| DB2725 | 141x *tetO* array @ *lac* (+1.10 Mb), Δ*recB745*::FRT*kan*FRT, pDM15 | This study |
| DB2954 | 141x *tetO* array @ *gln* (+0.13 Mb), *λc*I*ts857* $P_R$::*ruvC*Def-*gfp*::FRT*kan*FRT, pDM15 | This study |
| DB2956 | 141x *tetO* array @ *dnaB* (+0.34 Mb), *λc*I*ts857* $P_R$::*ruvC*Def-*gfp*::FRT*kan*FRT, pDM15 | This study |
| DB3007 | MG1655 *ssb-ypet*::*kan* | This study |
| DB3124 | 141x *tetO* array @ *lac* (+1.10 Mb), *λc*I*ts857* $P_R$::*ruvC*Def-*gfp*::FRT*kan*FRT, pDM15 | This study |
| DB3195 | 141x *tetO* array @ *ytjB* (+0.63 Mb), pDM15 | This study |
| DB3198 | 141x *tetO* array @ *alsC* (+0.32 Mb), pDM15 | This study |
| DB3216 | 141x *tetO* array @ *yhgN* (−0.35 Mb), pDM15 | This study |
| DB3264 | 141x *tetO* array @ *yjfY* (+0.58 Mb), pDM15 | This study |
| JW2788 | Δ*recB*::FRT*kan*FRT | (Baba *et al.*, 2006) |
| RRL190 | AB1157 FRT*kan*FRT::*dnaN-ypet* | (Reyes-Lamothe R *et al.*, 2010) |
| RRL32 | AB1157 *ssb-ypet*::*kan* | (Reyes-Lamothe R. *et al.*, 2008) |
| SMR22529 | *λc*I*ts857* $P_R$::*ruvC*Def-*gfp*::FRT*kan*FRT | This study |
| pCP20 | Flippase FLP $Ap^R$ recombinase expression plasmid | (Cherepanov & Wackernagel, 1995) |
| pJZ087 | FRT-141x *tetO* $Gm^R$ integration plasmid | (Wang M *et al.*, 2019) |
| pDM15 | $P_{nahG}$::*tetR-yfp* $Cm^R$ expression plasmid | (Magnan & Bates, 2015) |

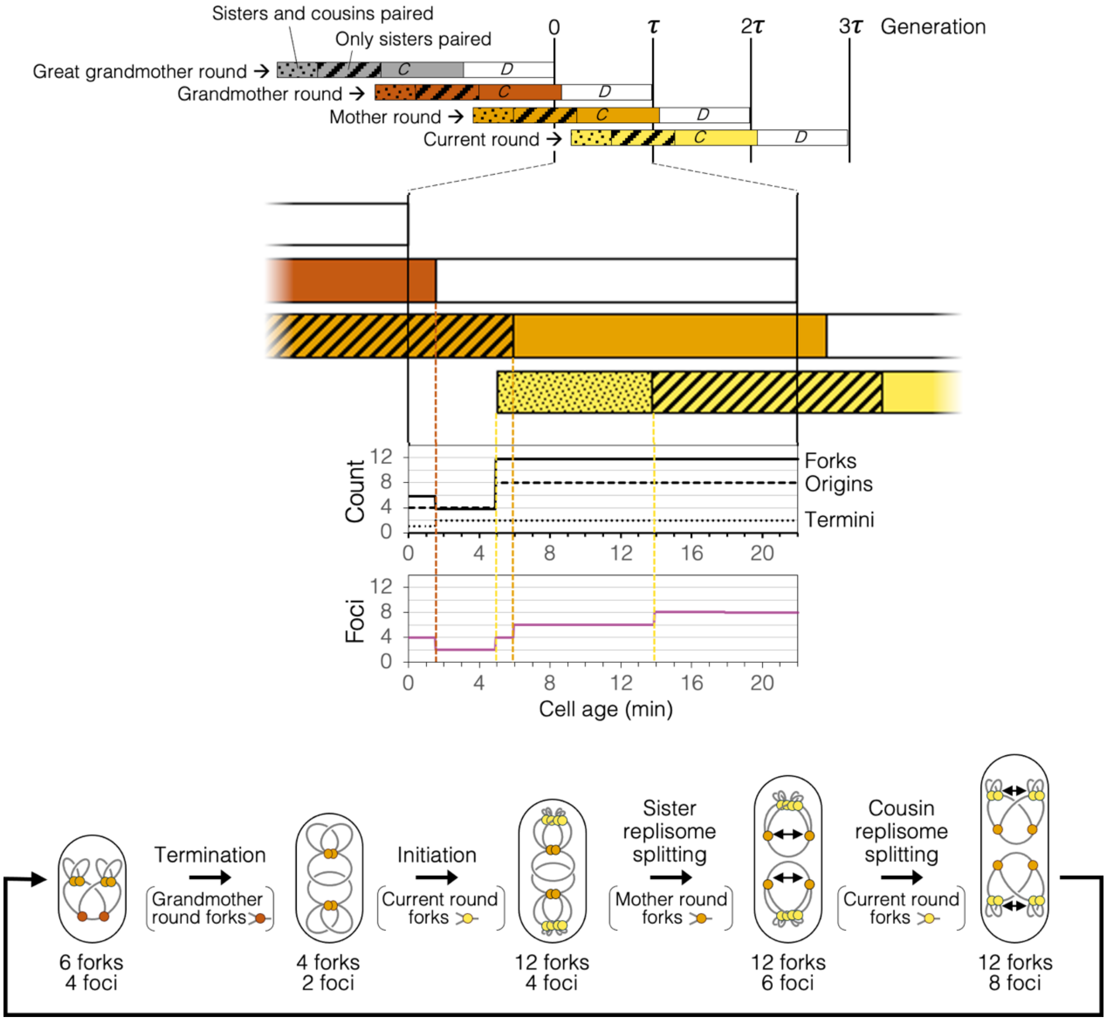


**Figure 2 – figure supplement 1.** Replication timeline and spatial dynamics under the Splitting with cousins model. Arrangement of replisomes and DNA in cell drawings are speculative and are based solely on measurements of genomic content and number of DnaN-YPet foci per cell (Figure 2H, I).

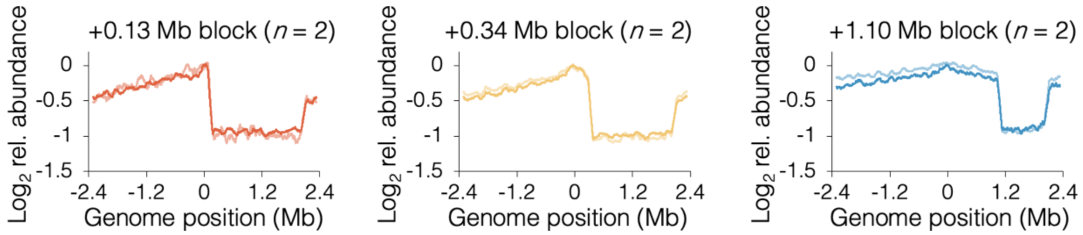


**Figure 3 – figure supplement 1.** 6 raw sequencing profiles of roadblocked cells under slow growth conditions. Profiles along the blocked chromosome arm are ~flat between the *tetO* array and the most distal *ter* site, which terminates replication emanating from the unblocked arm. Profile noise is a consequence of low copy number under slow growth conditions (see Figure 3 – figure supplement 3 for comparison).

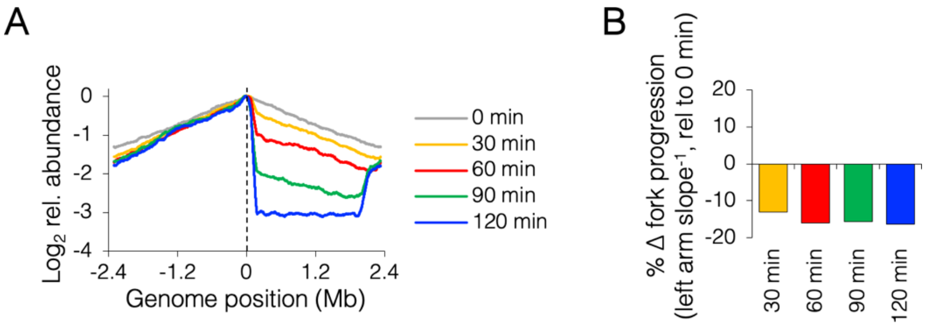


**Figure 3 – figure supplement 2.** Roadblocking time course under fast growth conditions. (A) Sequencing profiles for +0.13 Mb roadblock every 30 minutes after TetR-YFP induction (*n*=1). (B) Change in progression of the unblocked (left) replisome after roadblocking relative to time zero.

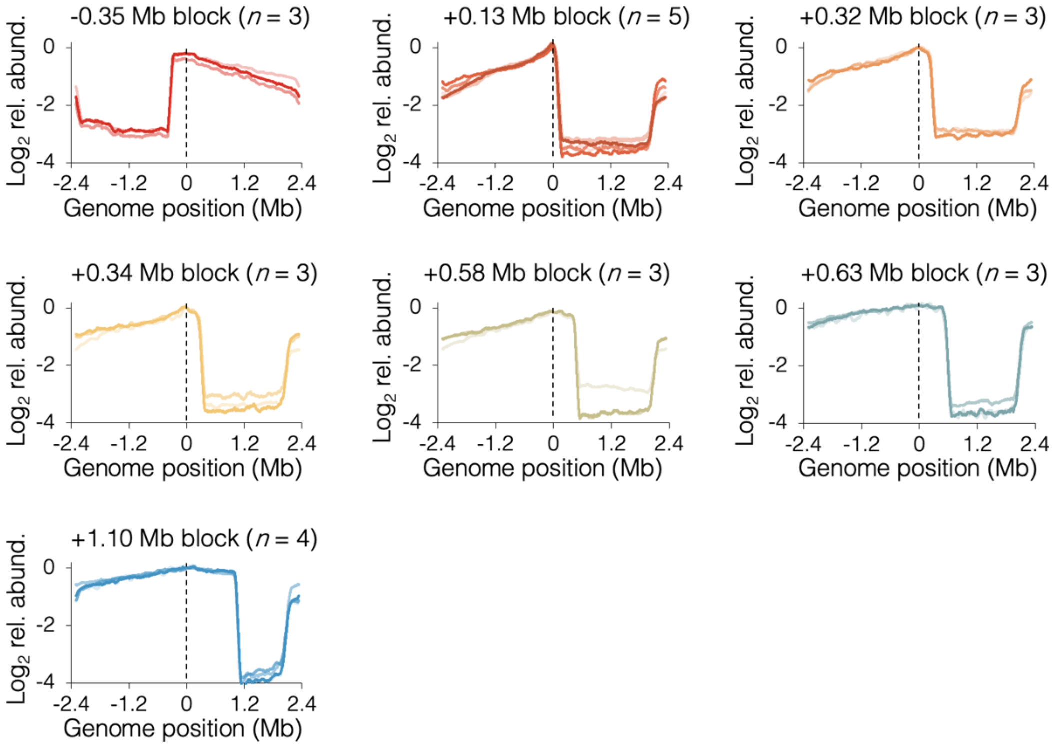


**Figure 3 – figure supplement 3.** 24 raw sequencing profiles of roadblocked cells under fast growth conditions. Profiles along the blocked chromosome arm are ~flat between the *tetO* array and the most distal *ter* site, which terminates replication emanating from the unblocked arm.

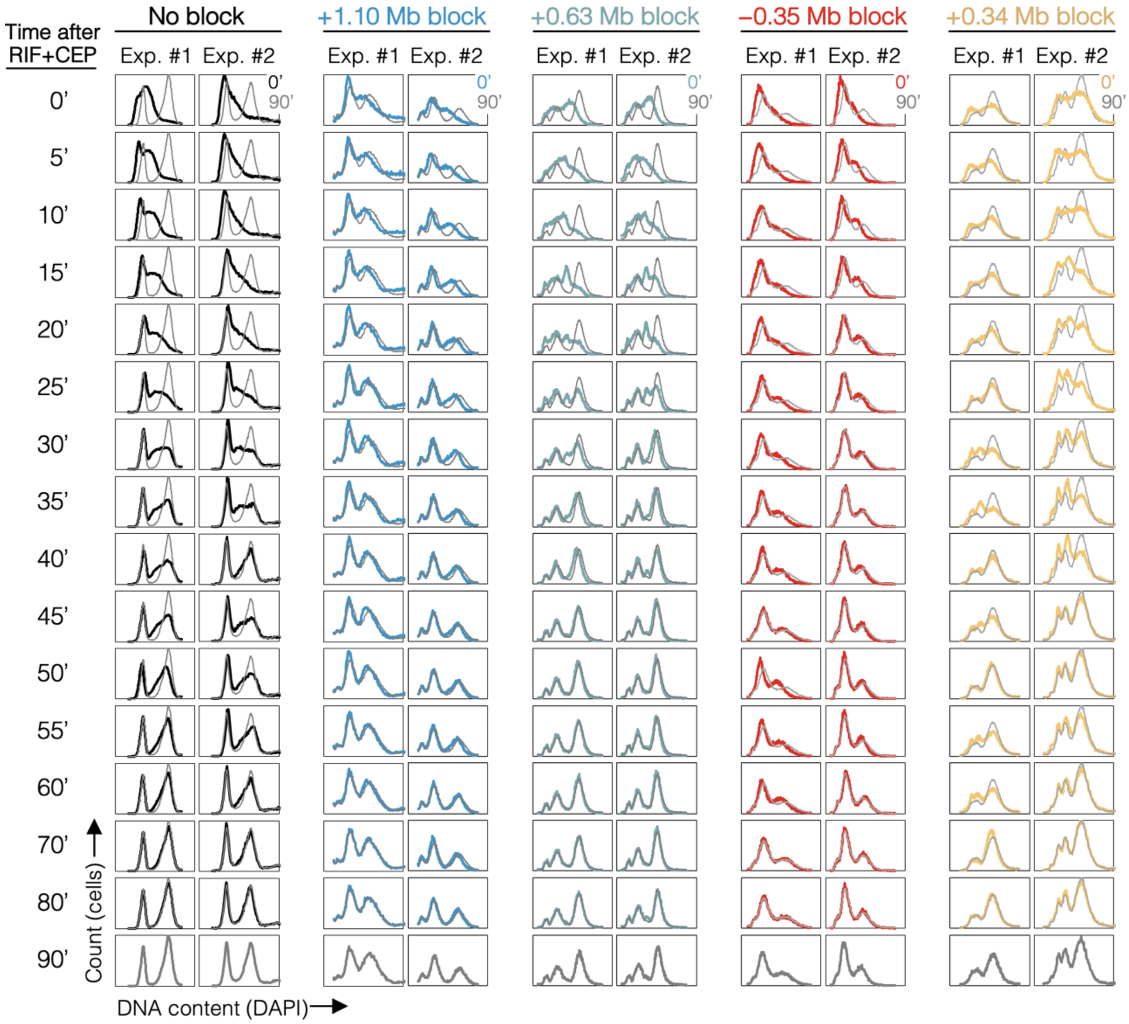


**Figure 4 – figure supplement 1.** Complete time courses for 10 rifampicin runout experiments. DNA histograms are shown at 5-minute intervals with 90 minute time point overlayed (grey).

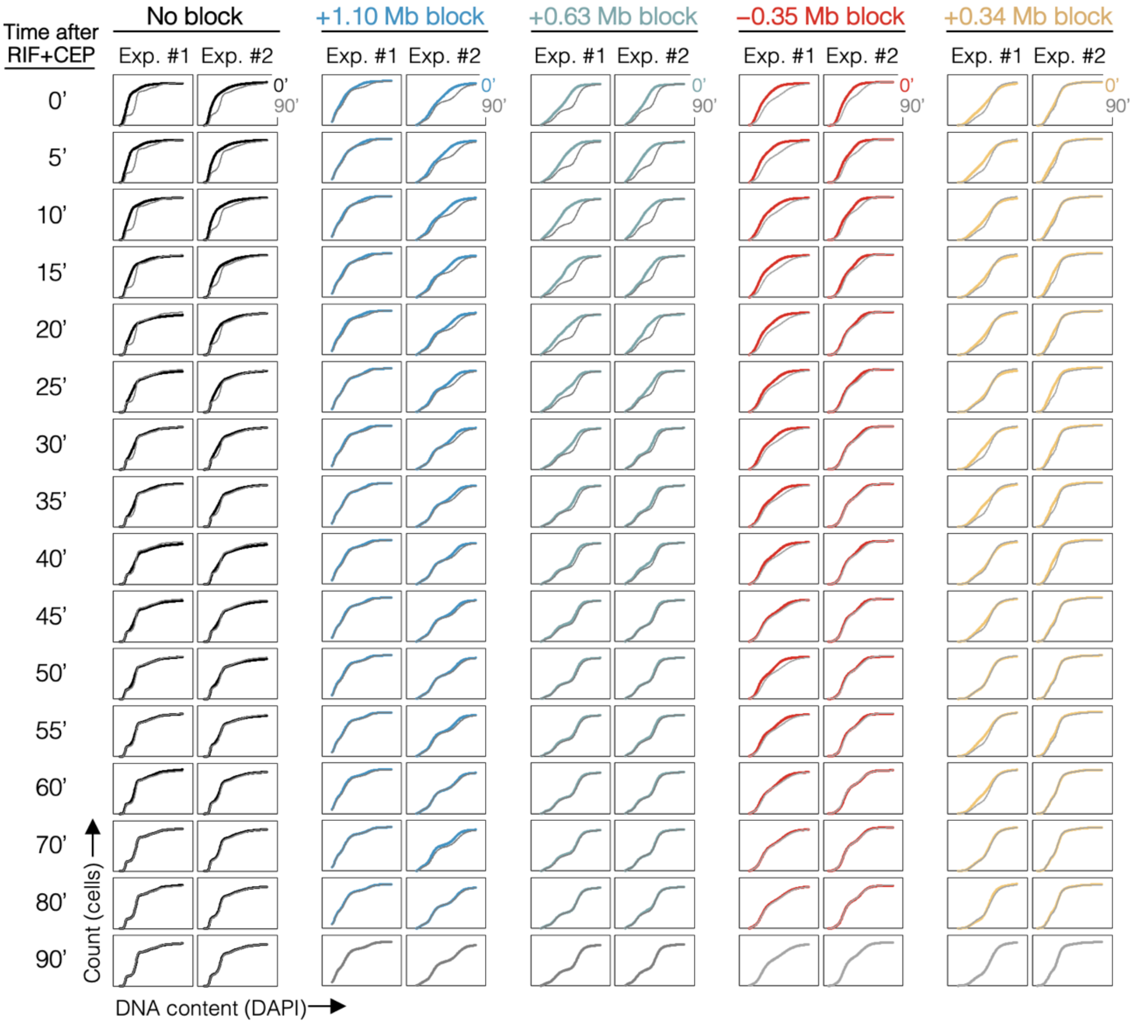


**Figure 4 – figure supplement 2.** Cumulative curve plots for 10 rifampicin runout experiments. Cumulative curves of DNA histograms (Figure 4 – figure supplement 1) with 90 minute time point overlayed (grey).

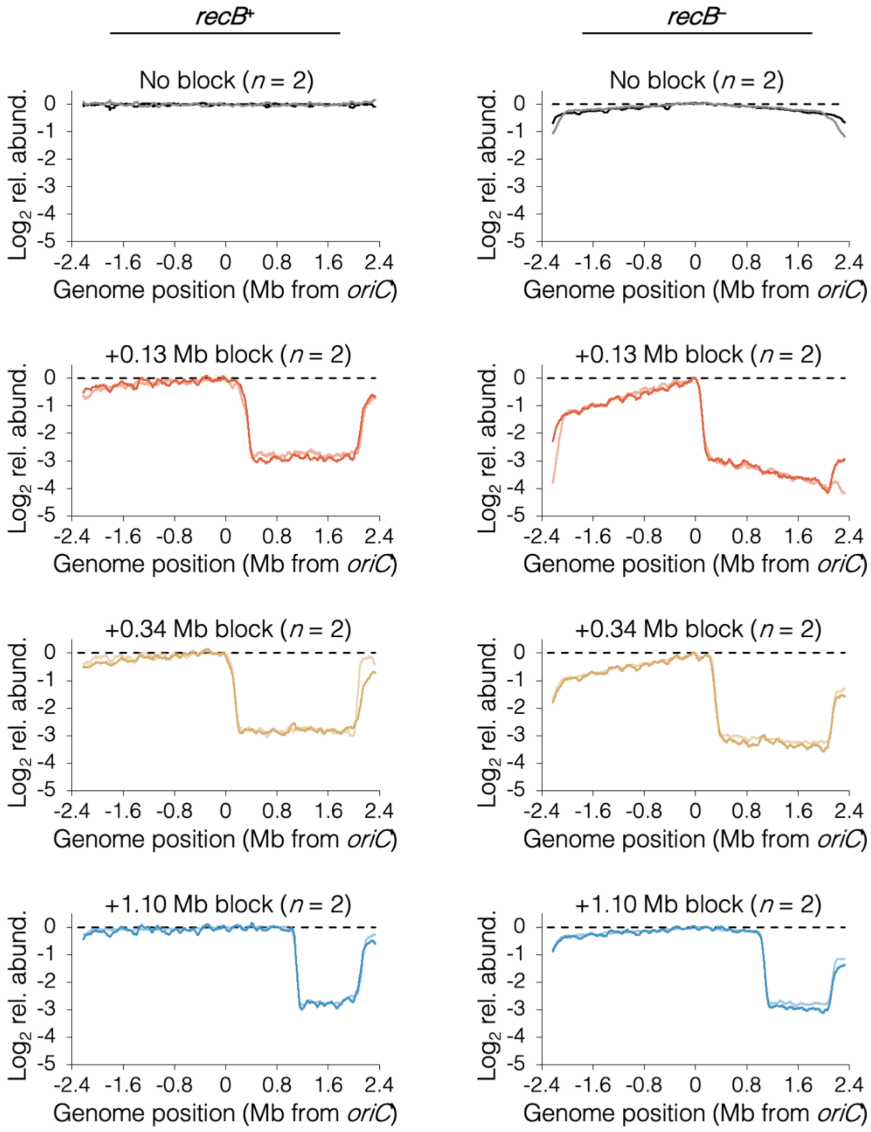


**Figure 5 – figure supplement 1.** 16 raw sequencing profiles of cells after rifampicin runout.

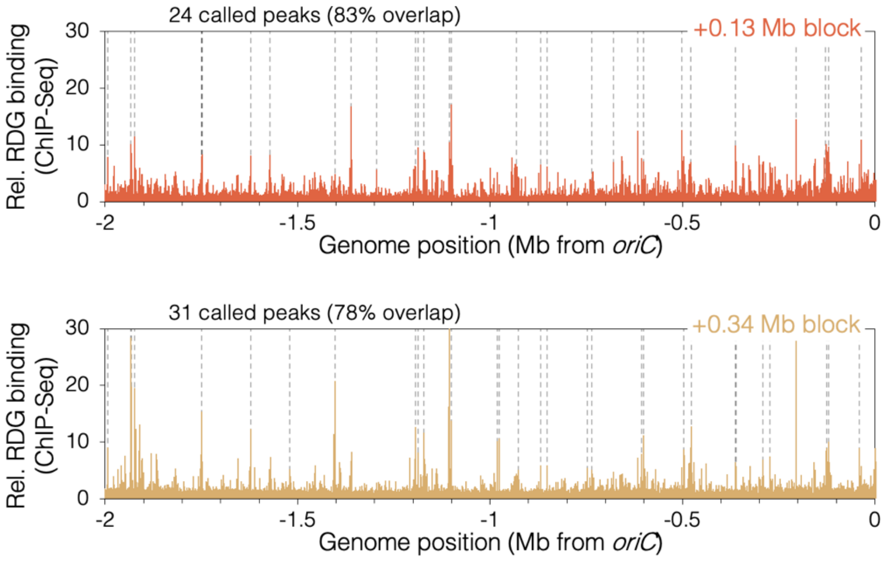


**Figure 6 – figure supplement 1.** RDG binding peaks in cells with a roadblock at +0.13 Mb or +0.34 Mb. Unblocked chromosome arm is shown between *oriC* and -2 Mb. Peaks (dashed lines) are locations of RDG signal ≥ 4 standard deviations above the 100-kb local median.

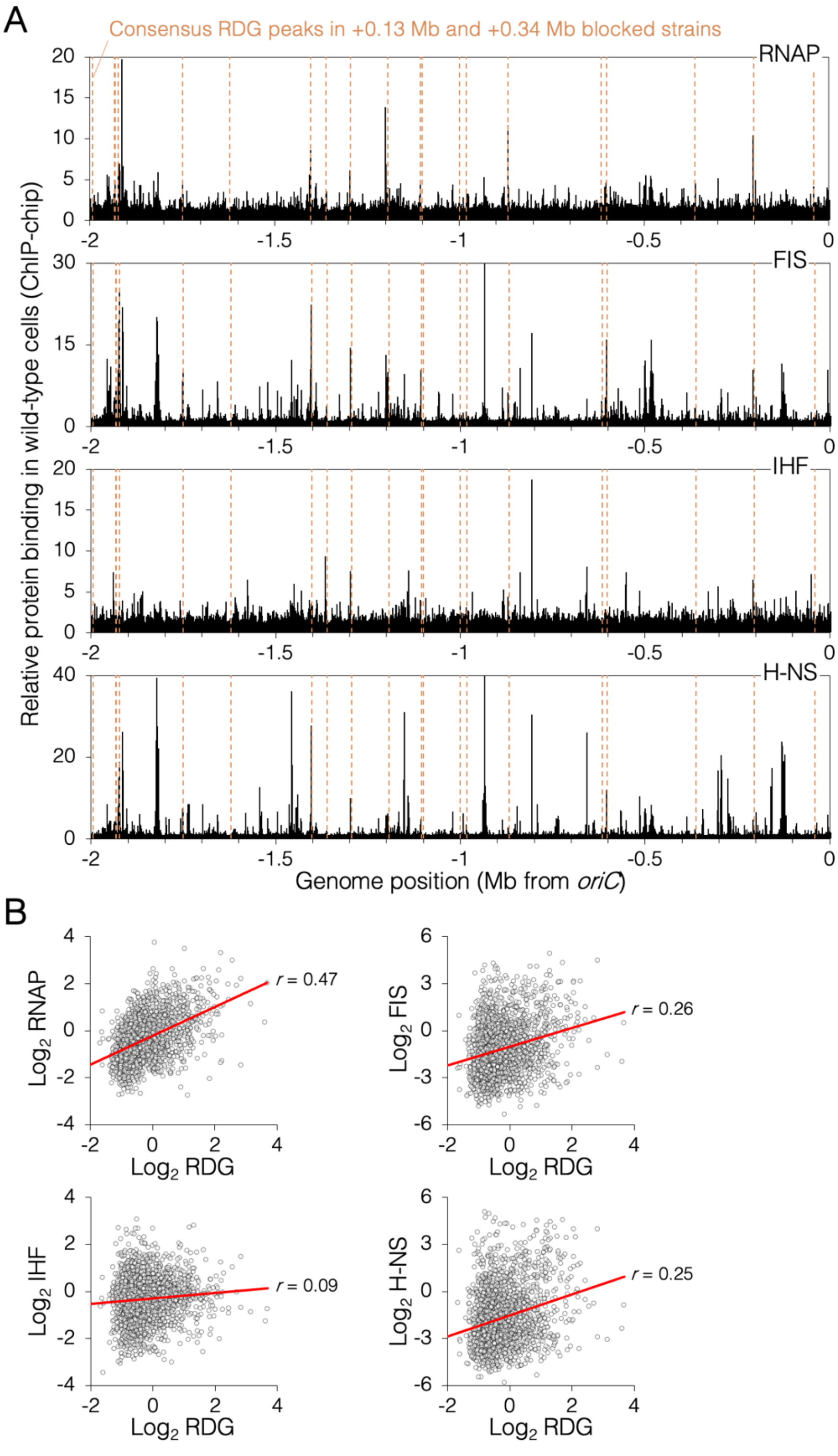


**Figure 6 – figure supplement 2.** Genomic binding of RDG and major nucleoid proteins. (A) Nuceoid protein binding profiles (black) and consensus RDG binding peaks (dashed lines). (B) Correlation plots of nucleoid protein binding and RDG binding (average +0.13 Mb, +0.34 Mb blocked cells). All data is along the unblocked chromosome arm only, between *oriC* and -2 Mb.
